# An executive center for the intake of liquids

**DOI:** 10.1101/2021.07.02.450862

**Authors:** Bowen Dempsey, Selvee Sungeelee, Philip Bokiniec, Zoubida Chettouh, Séverine Diem, Sandra Autran, Evan R. Harrell, James F.A. Poulet, Carmen Birchmeier, Harry Carrey, Auguste Genovesio, Simon McMullan, Christo Goridis, Gilles Fortin, Jean-François Brunet

## Abstract

It has long been known that orofacial movements for feeding can be triggered, coordinated, and often rhythmically organized at the level of the brainstem, without input from higher centers. We uncover two nuclei that can organize the movements for ingesting fluids in mammals. These neuronal groups, defined by unique transcriptional codes and developmental origins, IRt^*Phox2b*^ and Peri5^*Atoh1*^, are located, respectively, in the intermediate reticular formation of the medulla and around the motor nucleus of the trigeminal nerve. They are premotor to all jaw-opening and tongue muscles. Stimulation of either, in awake animals, opens the jaw, while IRt^*Phox2b*^ alone also protracts the tongue. Moreover, stationary stimulation of IRt^*Phox2b*^ entrains a rhythmic alternation of tongue protraction and retraction, synchronized with jaw opening and closing, that mimics lapping. Finally, fiber photometric recordings show that IRt^*Phox2b*^ is active during volitional lapping. Our study identifies one of the long hypothesized subcortical nuclei underpinning a stereotyped feeding behavior.

## Introduction

The hindbrain (medulla and pons) is a sensory and motor center for the head and the autonomic (or visceral) nervous system. Large areas therein defy conventional cytoarchitectonic description and are subsumed under the label “reticular formation” (*1*). Over decades, the reticular formation has slowly emerged from “localizatory nihilism” (*2*), and regions defined by stereotaxy [e.g.(*3*)], or cell groups defined by their projections [e.g.(*4*)] have been implicated in a variety of behaviors. Notably, the reticular formation contains premotor neurons to orofacial or respiratory muscles, and rhythm generators for chewing, licking, whisking, breathing and sighing (*3*)(*5*)(*6*)(*7*)(*8*)(*9*). However, the parsing of the reticular formation into genetically defined neuronal groups, endowed with specific connectivity and roles, has only begun (*10*)(*11*)(*12*)(*13*)(*14*) and lags behind other parts of the brain, such as the cortex or the spinal cord.

Among the most specific markers of neuronal classes are transcription factors, in particular homeodomain proteins [e.g. (*15*)(*16*)]. *Phox2b* is one such gene, which marks (and specifies) a limited set of neurons in the peripheral nervous systems and the hindbrain. The *Phox2b* expression landscape is strikingly unified by physiology: most *Phox2b* neurons partake in the sensorimotor reflexes of the autonomic nervous system, that control bodily homeostasis (*17*). An apparent exception are branchial motor neurons, that motorize the face and neck (*1*),(*18*) but their kinship to visceral circuits, aptly highlighted by their alternative name of “special visceral”, is revealed by their ancestral functions in aquatic vertebrates, exclusively in feeding and breathing — thus “visceral” indeed. Finally, *Phox2b* labels neurons in the reticular formation, with unknown functions so far, that we set out to explore in this study.

## Results

### The reticular formation harbors *Phox2b*^+^ orofacial premotor neurons

We visualized the total projections of *Phox2b* interneurons that are located in the reticular formation. To exclude the potentially confounding widespread projections of the noradrenergic locus coeruleus, which also expresses *Phox2b* (*19*), we exploited the glutamatergic nature of most reticular *Phox2b+* interneurons — thus their expression of the glutamate vesicular transporter *vGlut2* (**Fig. S1A**) — and a novel intersectional allele (*Rosa^FRTtomato-loxSypGFP^* or *Rosa^FTLG^*) (**Fig. 1A**) which expresses one of two fluorophores, exclusively: a cytoplasmic tdTomato (tdT) upon action of flippase (*FLPo*) or, upon additional action of *Cre* recombinase, a fusion of synaptophysin with GFP (*Syp-GFP*) transported to pre-synaptic sites (*20*). In the hindbrain of *Phox2b::Flpo;vGlut2::Cre;Rosa^FTLG^* pups at P4, tdT was expressed, as expected, in the singly recombined soma of the cholinergic *Phox2b^+^* branchiomotor (and visceromotor) neurons, but lost from the doubly recombined *vGlut2^+^ Phox2b^+^* interneurons (**Fig. S1B**), whose *Syp-GFP^+^* boutons covered discrete structures of the hindbrain (**Fig. S1B, Fig. 1B**). Among the targeted structures, motor nuclei featured prominently: *i)* most branchiomotor (*Phox2b*^+^) nuclei — the trigeminal motor nucleus (Mo5) and its accessory nucleus (Acc5), the facial nucleus (Mo7) (albeit only its intermediate lobe) and its accessory nucleus (Acc7), the nucleus ambiguus (MoA); *ii)* two somatic (*Phox2b*^—^) motor nuclei: the hypoglossal nucleus (Mo12), and a nucleus in the medial ventral horn, at the spinal-medullary junction, which innervates the infrahyoid muscles (*21*) (and **Fig. S1C)**, and that we call MoC (to denote its projection through the upper Cervical nerves) (*21*). Other cranial motor nuclei were free of input from *Phox2b^+^*/*vGlut2^+^* interneurons: those for extrinsic muscles of the eye (oculomotor (Mo3) and trochlear (Mo4)), and for the spinal accessory nucleus (Mo11), which innervates the sternocleidomastoid and trapezius muscles (**Fig. S1D**). The abducens nucleus (Mo6) however, did receive boutons (**Fig. S1D**). Thus, somewhere in the reticular formation are *Phox2b*^+^ orofacial premotor neurons, which we then sought to locate.

**Fig. 1.**
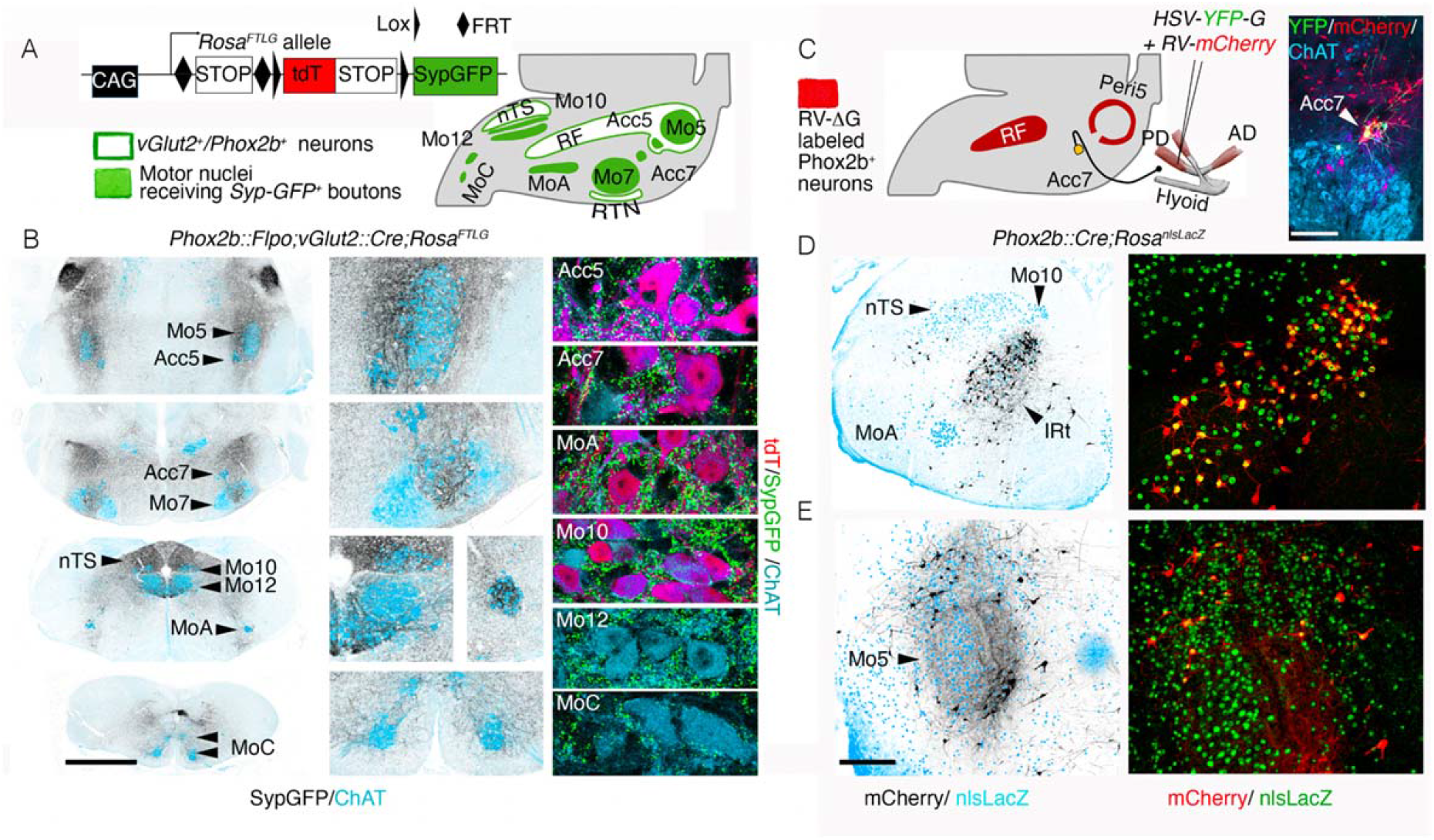
Premotor status of reticular formation Phox2b+ interneurons. (**A**) *Rosa^FTLG^* allele used for intersectional transgenic labeling of boutons from *vGlut2/Phox2b* interneurons (left) and schematic of the results (right). (**B**) Coronal sections through the hindbrain of a *Phox2b::Flpo; vGlut2::Cre;Rosa^FTLG^* mouse at P4, showing synaptic boutons (black) from *vGlut2/Phox2b* interneurons in relation to motor nuclei (ChAT^+^, blue) at low (left), and higher (middle) magnifications, and close ups of boutons (green) on motoneurons (right), which either *Phox2b*^+^ (purple) or *Phox2b*^—^ (blue). (**C**) (left) Strategy for mono-synaptically restricted trans-synaptic labeling of premotor neurons from the posterior digastric muscle (PD) in a *Phox2b::Cre;Rosa^nlsLacZ^* mouse, with G-deleted rabies virus (RV) encoding *mCherry* and complemented by a G-encoding helper HSV virus (*HSV-G-YFP*), and summary of the results. (right panel) The only seed neurons are Acc7 motoneurons, double-labeled by the *HSV-G* and *RV-mCherry* viruses. (**D**,**E**) Coronal sections through the hindbrain at P8 showing labeled premotor neurons (black on the left panels) in the medial IRt (**D**) and Peri5 (**E**), which for the most part (72.7%±3.5 SEM, n=4 animals) express *Phox2b* (right panels). AD, anterior digastric; IRt, intermediate reticular formation; nTS, nucleus of the solitary tract; PD, posterior digastric; peri5, peri-trigeminal area; RF, reticular formation; RTN, retrotrapezoid nucleus Scale bars, (**B**) 1mm for the left column, (**C**) 250μm, (**D**,**E**), 500μm.

To locate *Phox2b*^+^ orofacial premotor neurons, we used retrograde trans-synaptic viral tracing from oromotor muscles. We injected a G-defective rabies virus variant encoding *m-Cherry* (*22*) together with a helper virus (*HSV-G*) in the posterior belly of the digastric muscle (**Fig. 1C**) (a jaw-abductor), which is innervated by Acc7 (*23*)(*24*). Acc7 predictably contained the only seed neurons of the central nervous system (**Fig. 1C**), while *Phox2b^+^* premotor neurons were found at two sites: *i)* the intermediate reticular formation (IRt) (**Fig. 1D**), and *ii)* “regio h”, arranged in “shell form” around Mo5 (*25*), more commonly called the peritrigeminal region (Peri5) (*26*) (**Fig. 1E**). We found the same pattern of *Phox2b*^+^ premotor neurons for the geniohyoid muscle (a tongue protractor) (**Fig. S2A**), innervated by the accessory compartment of Mo12 (Acc12)(*21*)); and we found a subset of this pattern for the genioglossus (a tongue protractor and/or jaw abductor) (**Fig. S2B**) and for the intrinsic muscles of the tongue (**Fig. S2C**) (both innervated by Mo12), whereby *Phox2b*^+^premotor neurons were restricted to the IRt. On the other hand, the masseter (the main jaw closing muscle) and the thyro-arytenoid (that motorizes the vocal cords) had totally distinct premotor landscapes (**Fig. S2D,E**) (*27*)(*28*).

We next sought to characterize genetically and developmentally the *Phox2b*^+^ orofacial premotor neurons located in Peri5 and IRt.

### Transcriptional signature and developmental origin of Peri5^Atoh1^ and IRt^Phox2b^

The *Phox2b*^+^ premotor nucleus that occupies Peri5, we shall call Peri5^*Phox2b*^ (**Fig. 2A,B**). Because it surrounds, shell-like, a nucleus with a history of *Phox2b* expression —Mo5+Acc5 — it cannot be selectively accessed with *Phox2b*-based tools, even refined by stereotaxy. We thus restricted our study to a distinct subnucleus of Peri5^*Phox2b*^, which unlike the rest of the nucleus co-expresses *Phox2b* with another transcription factor, *Atoh1* (*29*) and that we shall call Peri5^*Atoh1*^ (**Fig. 2B-D**). Peri5^*Atoh1*^ is made of 2052±184 cells (n=4) at late gestation (E18.5), is premotor to the posterior digastric (**Fig. S2F**), and can be selectively targeted in an intersectional *Phox2b::flpo;Atoh1::Cre* background (**Fig. 2E**). Peri5^*Atoh1*^ cells express *Lbx1*(**Fig. 2F**), thus originate from the dB progenitor domain (*30*). More precisely they belong to its dB2 derivatives, at the leading edge of whose migration stream they become detectable at E11.5, near the incipient Mo5 (**Fig. 2G**).

**Fig. 2.**
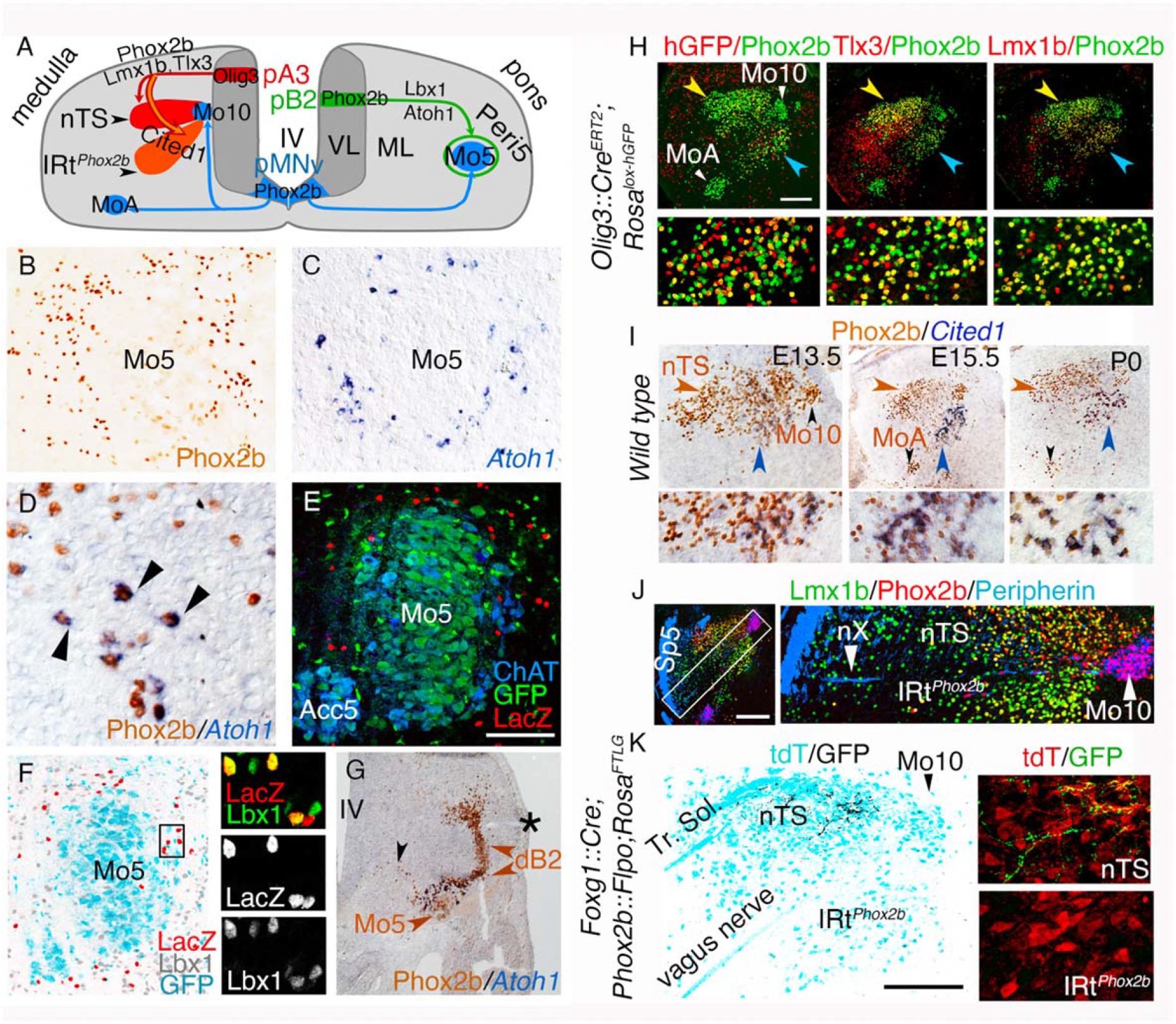
Ontogenetic definition of IRt^*Phox2b*^ and Peri5^*Atoh1*^. (**A**) Two schematic hemi-sections of the embryonic medulla (left) or pons (right), showing the origin of branchiomotor nuclei (Mo5, MoA and Mo10), Peri5^*Phox2b*^ and IRt^*Phox2b*^ in progenitor (p) domains of the ventricular layer (VL), their settling sites in the mantle layer (ML), and their transcriptional codes. (**B**,**C**,**D**) Coronal sections through the pons at E18.5, showing Peri5^*Phox2b*^ (**B**) or Peri5^*Atoh1*^ (**C**,**D**) labeled with the indicated antibody or probe. Peri5^*Atoh1*^ cells co-express *Phox2b* and *Atoh1* (arrowheads in **D**). (**E**) Coronal sections through the pons of a *Phox2b::Flpo;Atoh1::Cre;Fela* mouse at P0, showing the double-recombined (nlsLacZ^+^) cells of Peri5^*Atoh1*^ (red). (**F**) Coronal section through Mo5 in a *Phox2b::Flpo;Atoh1::Cre;Fela* mouse, where *Phox2b^+^* motoneurons are *GFP^+^* (cyan) and *Phox2b^+^/Atoh1^+^* neurons are *nlsLacZ^+^* (red), counterstained for *Lbx1* (grey at low magnification, green in the close ups). (**G**) Coronal section through the pons at E11.5 showing the migrating *Phox2b^+^* Mo5 and dB2 precursors (black and brown arrowheads respectively) and, at their meeting point, Peri5^*Atoh1*^ cells that have switched on *Atoh1*. Asterisk: lateral recess of the IVth ventricle (IV). (**H**) Coronal sections through nTS (yellow arrowhead) and IRt^*Phox2b*^ (blue arrowhead) at E18.5, at low (upper) or high (lower) magnification, stained with the indicated antibodies. A history of *Olig3* expression is revealed by recombination of the histone-GFP (hGFP) reporter in the *Olig3::Cre^ERT2^* background (left). Mosaicism is likely due to incomplete induction of Cre. Virtually all cells of IRt^*Phox2b*^ (98% ± 0.2 SEM, n=3 animals) co-expressed *Lmx1b* with *Phox2b*. (**I**) Coronal sections through nTS (brown arrowhead) and IRt^*Phox2b*^ (blue arrowhead) at indicated stages at low (upper) add high (lower) magnification, immunostained for *Phox2b* and in situ hybridized for *Cited1*. (**J**) Coronal section at E15.5 showing that nTS and IRt^*Phox2b*^ are separated by the medullary root of the vagus nerve (nX). Sp5, spinal trigeminal tract. (**K**) Coronal section through the nTS and IRt^*Phox2b*^ of an adult, showing the central boutons of epibranchial ganglia (that express *Foxg1* (*68*) and are labeled by *SypGFP* in a *Foxg1^iresCre^;Phox2bFlpo;Rosa^FTLG^* background) in the nTS, but not IRt^*Phox2b*^ (left). Magnified details (right). Scale bars, (**E**) 100 μm; (**H, J, K**) 200 μm.

The *Phox2b^+^* premotor nucleus that occupies IRt, we shall call IRt^*Phox2b*^ (**Fig. 2A**). It shares with the nearby nTS the *Phox2b/Tlx3/Lmx1b* signature and an origin in *Olig3^+^* progenitors (i.e. the pA3 progenitor domain (*31*)) (**Fig. 2A,H**). It is distinguished, however, by expression of the transcriptional cofactor *Cited1* (**Fig. 2I**). IRt^*Phox2b*^ segregates topographically from nTS at E13.5 (**Fig. 2I**) from which it can thus be told apart by stereotaxy. The border between the two nuclei is marked by the intramedullary root of Mo10 (**Fig. 2J**). Unlike nTS, IRt^*Phox2b*^ does not receive any input from the tractus solitarius (**Fig. 2K**). Thus, IRt^*Phox2b*^ and nTS are two structures related by lineage, which acquire distinct molecular, topological and hodological identities.

### Peri5^*Atoh1*^ and IRt^*Phox2b*^ target jaw opening and tongue muscles

We confirmed the premotor status of Peri5^*Atoh1*^ or IRt^*Phox2b*^ in adult animals by anterograde tracing with viral and transgenic tools (**Fig. 3**). For Peri5^*Atoh1*^, we used the *Rosa^FTLG^* allele recombined by *Phox2b::Flpo* (*32*) and *Atoh1::Cre* (*13*) (**Fig. 3A**). The *GFP^+^* boutons covered Acc5, intermediate Mo7, Acc7, Mo10, Mo12 and MoC (**Fig. 3A-F**). In Mo12, the rostro-ventral compartment was excluded **(Fig. 3D,E**). Because the retrotrapezoid nucleus (RTN) is also *Atoh1^+^/Phox2b^+^*(*13*), thus could confound this pattern, we confirmed the projections of Peri5^*Atoh1*^ by anterograde tracing with a *Cre*-dependent adeno-associated virus (AAV) expressing *mGFP* and *Syp-mRuby* (*33*) injected in Mo5 of a mouse harboring both, *Phox2b-Flpo* and an *Atoh1-Cre* that is dependent on *Flpo* (*Atoh1::FRTCre*)(*13*) (**Fig. S3A,B**). Using the same vector stereotaxically injected in IRt^*Phox2b*^ of a *Phox2b::Cre* mouse, we found the projections from IRt^*Phox2b*^ in the same motor nuclei as those from Peri5^*Atoh1*^ (**Fig. 3G-L**)— with the notable difference that in Mo12, the ventral compartment was targeted, rather than the dorsal one (compare **Fig. 3J,K** with **Fig. 3D,E**).

**Fig. 3.**
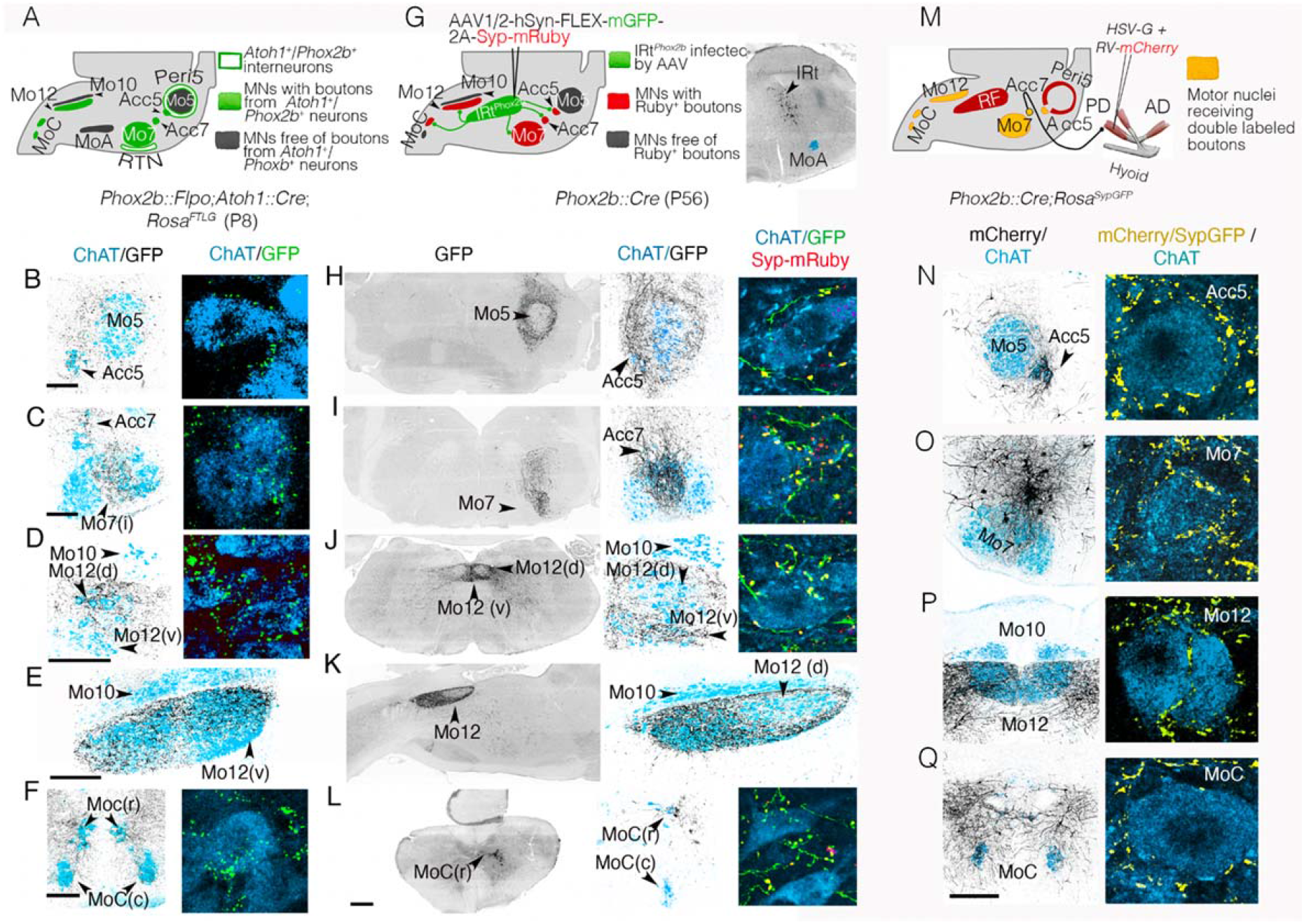
Projections of IRt^*Phox2b*^ and Peri5^*Atoh1*^ on hindbrain motoneurons. (**A**) Strategy for the transgenic labeling of projections from Peri5^*Atoh1*^ (and RTN) and summary of the results; (**B**-**F**) Coronal (**B**-**D**,**F**) or parasagittal (**E**) sections through a P8 hindbrain showing GFP-labeled boutons (black) on motoneurons (blue) at medium (left) and high (right) magnification. (**G**) Strategy for the viral tracing of projections from IRt^*Phox2b*^ and summary of the results (left), and mGFP-labeled infected cells of IRt^*Phox2b*^ (right); (**H**-**L**) Coronal (**H**-**J**,**L**) or parasagittal (**K**) sections through a P56 hindbrain showing the GFP-labeled fibers (black) of IRt^*Phox2b*^ neurons at low (left) and medium (middle) magnifications, and in extreme close-ups (right), together with Syp-mRuby labeled boutons on motoneurons (blue). Scale bar, (**B-F**) 200 μm, (**H-L**) 500 μm. (**M**) schematic for retrograde tracing of premotor neurons for the right posterior digastric muscle, in a *Phox2b::Cre;Rosa^SypGFP^*, and summary of the results. (**N**-**Q**) (left) Coronal sections through the hindbrain at P8 showing the *mCherry*^+^ projections (black) of premotor neurons on the motor nuclei (*ChAT*^+^,blue); (right) close ups on motoneurons receiving double-labeled *Spy-GFP*/*mCherry* boutons (yellow). Scale bars, (**B-F**) 200μm for the left column, (**H-L**) 500 μm for the left column, (**N-Q**) 200 μm for the left column.

To map putative collaterals of *Phox2b^+^* premotor neurons, we performed a retrograde transsynaptic tracing experiment from the posterior digastric in a genetic background that, in addition, labels the boutons of all *Phox2b^+^* neurons with *GFP* (*Phox2b::Cre;Rosa::Syp-GFP*) (**Fig. 3M**). Double-labeled terminals (*m-Cherry^+^;Syp-GFP^+^*) — thus, sent by neurons that are both, *Phox2b^+^* and premotor to the posterior digastric — were found, in addition to Acc7 (the motor nucleus of the injected muscle), in Acc5, intermediate Mo7, Mo12 and MoC (**Fig. 3N-Q**). Thus, *Phox2b^+^* orofacial premotor neurons to Acc7 are collateralized in a way that hardwires Acc5, intermediate Mo7, Acc7, Mo12 and MoC to activate their target muscles together.

The combined action of head motor nuclei innervated by Peri5^*Atoh1*^ and IRt^*Phox2b*^ should mobilize the jaw, lower lip and tongue: Acc5 and Acc7 innervate the four suprahyoid muscles(*34*)^,^(*35*)^,^(*36*), which depress the jaw via the hyoid apparatus. Intermediate Mo7 innervates the *platysma* (*36*), probably a jaw depressor (*37*), and a *mentalis* (*36*), which, together with the *platysma*, pulls down the lower lip. Ventral Mo12, targeted by IRt^*Phox2b*^, innervates tongue protractors (*38*), while dorsal Mo12, targeted by Peri5^*Atoh1*^, innervates tongue retractors (*39*). Finally, MoC innervates the infrahyoid muscles, classically viewed as stabilizers of the hyoid during jaw lowering, but which probably collaborate with the suprahyoids in a more complex fashion (*40*).

In addition to motor nuclei, IRt^*Phox2b*^ projected massively to the peri5 region (**Fig. 3H**) and Peri5^*Atoh1*^ projected massively to IRt (**Fig. S3C**), including IRt^*Phox2b*^ (**Fig. S3D**, inset). Thus, Peri5^*Atoh1*^ and IRt^*Phox2b*^ appear reciprocally connected and in a position to, collectively, lower the jaw, while retracting or protracting the tongue, respectively.

### Peri5^*Atoh1*^ and IRt^*Phox2b*^ can trigger tongue and jaw movements

We optogenetically stimulated IRt^*Phox2b*^ or Peri5^*Atoh1*^ in head-fixed awake animals. To do so, we injected a *Cre*-dependent AAV that directs expression of the *CoChR* opsin to the cell soma, either in IRt^*Phox2b*^ of *Phox2b::Cre* mice (**Fig. 4A**), or in Peri5^*Atoh1*^ of *Phox2b::Flpo*;*Atoh1^FRTCre^* mice (**Fig. 4B**). Single light pulses (50ms) on IRt^*Phox2b*^ evoked a wide opening of the mouth accompanied by tongue protraction, which terminated upon cessation of the pulse (**Fig. 4A**), while the same stimulus applied to Peri5^*Atoh1*^ triggered only mouth opening, of smaller amplitude (**Fig. 4B**). Thus, both nuclei can open the mouth, in agreement with their projections on the motoneurons for the suprahyoid and infrahyoid muscles (**Fig. 1B, Fig. S2A, Fig. 3**), while IRt^*Phox2b*^ but not Peri5^*Atoh1*^ can protract the tongue, in line with the targeting of hypoglossal motoneurons for tongue protractors by the former and tongue retractors by the latter (**Fig. 3D,E,J,K**). Lengthening the light pulse on IRt^*Phox2b*^ (to 100ms or 200ms) analogically prolonged the mouth opening and tongue protraction (**Fig. 4C**). Unexpectedly however, further lengthening led to termination of the initial movement and its rhythmic repetition at around 7Hz (**Fig. 4C, Fig. S4A, Movie 1**), a frequency similar to that of naturally occurring licking (**Fig. S4B**) (*41*). Prolonged illumination of Peri5^*Atoh1*^ only prolonged the initial mouth opening (**Fig. 4D, Fig. S4C, Movie 2**). Thus, a contrast between the actions of photo-stimulated Peri5^*Atoh1*^ and IRt^*Phox2b*^ lies in the ability of the latter to translate stationary excitation into a rhythmic series of oromotor movements, akin to naturally occurring licking (*41*). This action requires that IRt^*Phox2b*^, on the one hand, engages a circuit that allows for delayed activation of antagonistic muscles. One such circuit might comprise the reciprocal projections of Peri5^*Atoh1*^ and IRt^*Phox2b*^ (**Fig. 3H, Fig. S3C,D**) for patterning the alternation of tongue protractions and retractions. In addition, IRt^*Phox2b*^ triggers rhythmicity and thus must be the long-hypothesized licking rhythm generator (*42*), or an element thereof — akin to another nearby *Phox2b^+^* nucleus, the RTN, which has rhythmic properties, in that case related to breathing, in the neonate (*43*)(*12*).

**Fig. 4.**
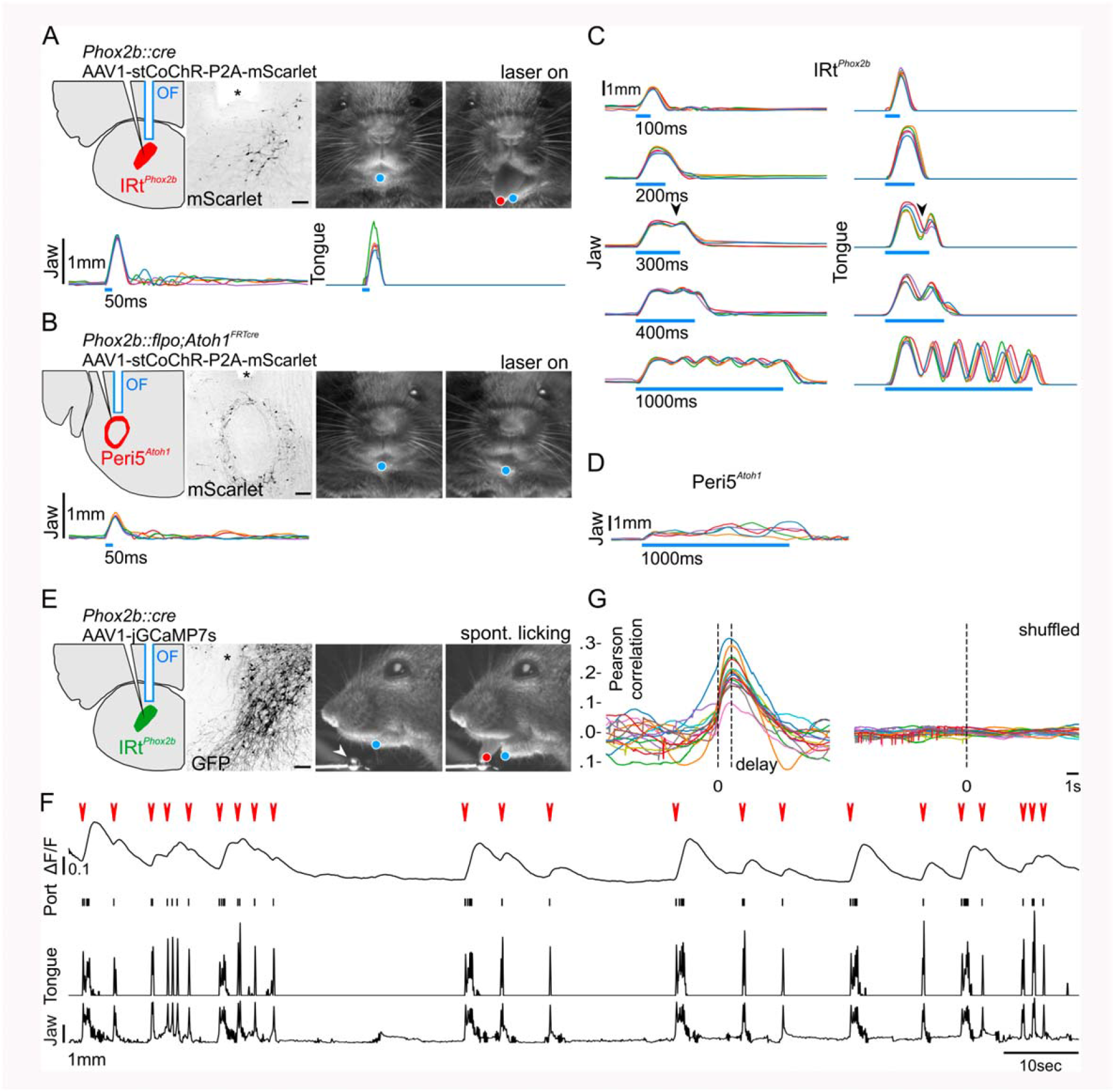
Orofacial movements triggered by IRt^*Phox2b*^ **and Peri5**^*Atoh1*^ **and activity of IRt**^*Phox2b*^ during voluntary licking. (**A**) (Upper left) Schematic of the viral injection and fiber optic implantation doe stimulation of IRt^*Phox2b*^ and transverse section though the hindbrain showing transduced IRt^*Phox2b*^ neurons and position of optical fiber (OF, asterisk); scale bar 100 μm. (Upper right) Example frames of the mouse face before and during stimulation including DeepLabCut tracked position of jaw (blue) and tongue (red). (Lower) Individual traces of tracked jaw and tongue position on the Y-axis upon 50 ms stimulation (5 trials). (**B**) (Upper left) Schematic of the viral injection and fiber optic implantation for stimulation of Peri5^*Atoh1*^ and transverse section though the hindbrain showing transduced Peri5^*Atoh1*^ neurons and position of optical fiber (asterisk); scale bar 200 μm. (Upper right) Example frames of the mouse face before and during stimulation including DeepLabCut tracked position of jaw (blue). (Lower) Individual traces (5 trials) of tracked jaw position on the Y-axis upon 50 ms stimulation. (**C**) Individual traces (5 trials) of tracked jaw (left) and tongue (right) position on the Y-axis upon stimulation of IRt^*Phox2b*^ of increasing length. A repetitive movement is triggered by stimulation beyond 300 ms (arrowhead). (**D**) Individual traces (5 trials) of tracked jaw position on the Y-axis upon a 1000 ms stimulation of Peri5^*Atoh1*^. The jaw remains open and quivers non-rhythmically during the stimulus. (**E**) (left) Schematic of viral injection and optical fiber implantation for observation of IRt^*Phox2b*^ activity, and transverse section through the hindbrain showing transduced IRt^*Phox2b*^ neurons and position of optical fiber (asterisk); scale bar 100 μm. (Middle) Example frames of the mouse face before and during a bout of licking from a lick port (arrowhead), during a photometry recording, including DeepLabCut tracked position of jaw (blue) and tongue (red). (**F**) Example trace of change in bulk fluorescence of IRt^*Phox2b*^ during a recording session (∼2 mins) of unitary licking events and licking bouts (red arrowheads), contact events with the lick port and movements of the tongue and the jaw on the Y axis. (**G**) (left) Superimposed correlation curves between licking activity and calcium activity (each curve corresponding to one of 15 recording sessions, each 1-5 min, in 1 mouse) which peaked at 1.2s after lick port contact; (right) no peak was observed after shuffling the data.

### IRt^Phox2b^ is active during volitional licking

We then tested whether IRt^*Phox2b*^ is active during spontaneous fluid ingestion. We recorded the bulk fluorescence (*44*) of IRt^*Phox2b*^ in head-fixed *Phox2b::Cre* mice, injected in IRt^*Phox2b*^ with a *Cre*-dependent AAV encoding the calcium indicator *jGCamp7s* (*45*) and implanted with an optical cannula (**Fig. 4E**). During freely initiated bouts of licking from a water-spout, we observed a systematic increase in fluorescence of IRt^*Phox2b*^ immediately upon deflection of the jaw that preceded individual licks or bouts of lapping (**Fig. 4F,G**, **Movie 3**). Thus, IRt^*Phox2b*^ neurons, capable of triggering a licking behavior with physiological frequency, are active during such spontaneous behavior. Importantly, IRt^*Phox2b*^ encompasses the location of many neurons identified as rhythmically active during licking (*6*). Stationary optogenetic stimulation of this nucleus might emulate the effect of sustained drive from the licking area of the oromotor cortex (*46*)(*47*)(*48*)(*49*).

### Inputs to IRt^*Phox2b*^

Although decerebrated mammals can display “reflexive” licking (*50*)(*51*), volitional or self-initiated licking requires higher brain centers. To explore the substratum for this requirement, we traced the inputs to IRt^*Phox2b*^ by co-injecting it with a pseudotyped G-defective rabies virus variant encoding *m-Cherry* and a helper virus that depends on *Cre*, in a *Phox2b::Cre* background (**Fig. 5A**). The vast majority of inputs (about 90%) were in the brainstem (**Fig. 5B**), which could explain the largely intact reflexive behavior of decerebrated animals. Among these regional inputs, many were found in IRt itself, including contralaterally (**Fig. 5C**) — suggesting local interconnectivity of IRt neurons, possibly related to rhythmogenesis. Other regional inputs came from the peri5 region (**Fig.5D**) — likely including Peri5^*Atoh1*^ that we had traced anterogradely to IRt^*Phox2b*^ (**Fig. S3D**)—, the mesencephalic nucleus of the trigeminal nerve (Mes5) (**Fig. 5E**) — which harbors proprioceptors for the teeth and masseter, potentially allowing for a cross-talk between jaw position and tongue movement (*53*), and the superior colliculi (**Fig. 5F**) —whose inhibition disrupts self-initiates licking (*52*). Finally, we found an input from the cortex (**Fig. 5G**), where a subclass of pyramidal tract neurons are known to directly target orofacial promotor neurons (*48*).

**Fig. 5.**
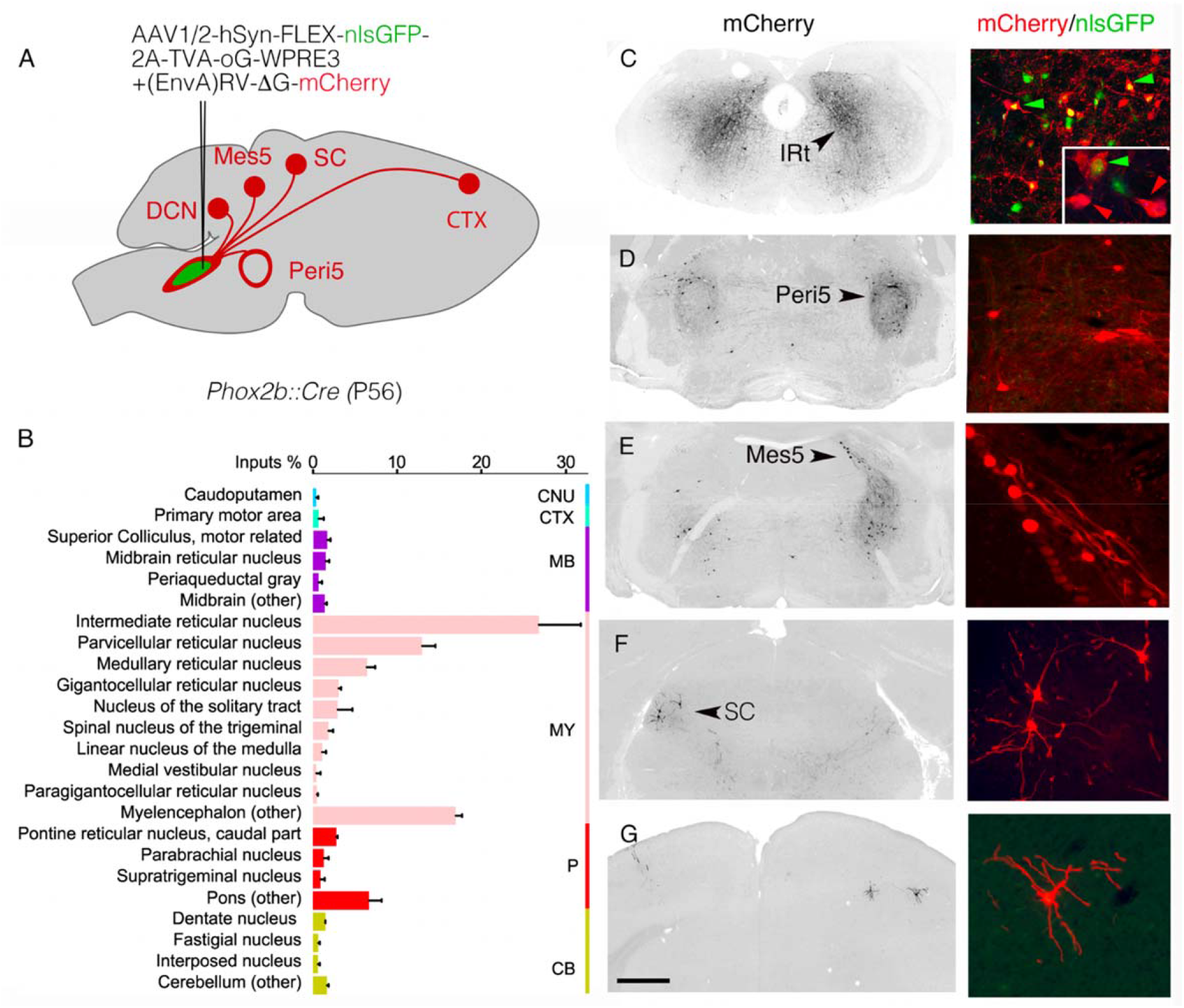
Inputs to IRt^Phox2b^. (**A**) Strategy for the retrograde trans-synaptic labeling of input neurons to IRt^*Phox2b*^ with exemplar sites of input. (**B**) Bar graph of the relative percentage of monosynaptic input neurons (1354±596 SEM) labelled from IRt^*Phox2b*^ starter neurons (199±91 SEM) from 3 animals (seeding efficiency 7.4±1.8 inputs/starter neuron), per brain region as defined in the Allen Brain Atlas. Rabies-labelled input neurons were largely (74.5±1.1%) restricted to the medulla (pink, MY) and exhibited a slight but consistent ipsilateral bias (55.6±3.0%). Major sources of these medullary inputs were the Intermediate, Gigantocellular and Parvocellular Reticular Nuclei. Inputs from the cortex, midbrain and pons represented a minority of rabies-labelled neurons (1.6±0.3%, 6.4±0.5% & 12.3±1.1% respectively). (**C**,**G**) Images at low magnification (left) and high magnification (right) of monosynaptic input neurons in the IRt, Peri5, mesencephalic nucleus of the trigeminal nerve, contralateral superior colliculus and motor cortex. Green arrowheads: seed neurons; red arrowheads: (n-1) IRt neurons. CB, cerebellum; CNU, caudoputamen; CTX, cortex; DCN, deep cerebellar nucleus; MB, midbrain; Mes5, mesencephalic nucleus of the trigeminal nerve; MY, myelencephalon; P, pons; SC: superior colliculus. Scale bar, 1mm.

## Discussion

Our study uncovers two genetically coded neuronal groups in the reticular formation, involved in orofacial movements. They are premotor to orofacial muscles and collaterized, thus in a position to coordinate the contraction of a precise set of muscles to the exclusion of others, a property previously highlighted in studies of orofacial premotor neurons ((*9*) (and references therein). As such, they represent an essential hierarchical level in the orchestration of complex oropharyngeal behaviors. In addition, one of them, IRt^*Phox2b*^, likely corresponds to the hypothetical licking CPG (*42*) which had been putatively located by correlative and lesioning studies ((*6*)(*5*) for reviews), or part thereof. So far, discussions of orofacial premotor neurons and rhythm generators have remained non-committal as to their mutual relationship (*6*)(*5*)(*9*). The most parsimonious interpretation of IRt^*Phox2b*^ is that its neurons are bifunctional: premotor through their collaterized inputs on motor nuclei, and rhythm generators. We cannot exclude at this stage, though, that IRt^*Phox2b*^ encompasses two subtypes of *Phox2b^+^* neurons, one premotor and the other pre-premotor, in charge of rhythm generation, and that it is the entrainment of the latter by photostimulation which triggers the rhythmic repetition.

IRt^*Phox2b*^ and Peri5^*Atoh1*^ as well as most of their motor targets, express the pan-autonomic *Phox2b* transcriptional determinant. Thus, the evolutionary conserved selectivity of *Phox2b* for neurons involved in homeostasis (*54*), extends beyond the reflex control of the viscera, including all sensory-motor loops involved in digestion (*17*)(*55*), to the executive control of ingestion, through the *Phox2b^+^* premotor/motor arm that mobilizes visceral-arch derived muscles. (**Fig. 1 and Fig. S1**). The remarkable genetic monotony of these circuits breaks down at the level of the somatic (*Phox2b^—^)* lingual and hypobranchials motoneurons. These exceptions are to be expected in the head where the visceral and somatic bodies of the vertebrate animal, *sensu* Romer (*56*), must meet and cooperate, at the border of the external world and interior milieu. Indeed, feeding can be construed as a sequence of somatic (i.e. external or relational) and visceral (i.e. internal or homeostatic) actions: to take in a substrate from the environment by biting or licking/lapping up, then to incorporate it in the interior milieu by chewing and swallowing. In these actions, the hyoid bone act as a weld between visceral and somatic muscles: the suprahyoids, derived from visceral arch mesoderm and innervated by branchiomotor (*Phox2b^+^*) motoneurons; and the “hypobranchials” (infrahyoid and lingual) derived from somites and innervated by somatic (*Phox2b*^—^) motoneurons. Hyoid bone, branchiomeric muscles, branchiomotor neurons and premotor centers Peri5^*Atoh1*^ and IRt^*Phox2b*^, all affiliated to the visceral body — muscles and bones through their origin in branchial arch mesoderm or neural crest, neurons through their expression of *Phox2b* — are likely the ancestral agents of feeding behaviors in vertebrates. At the advent of predatory and terrestrial lifestyles, the *Phox2b^+^* premotor centers must have recruited elements of the somatic body: the infrahyoid and lingual motoneurons, and their muscle targets, migrated into the head (*57*).

## Materials and Methods

### Mouse lines

The following transgenic mouse lines were used: *Phox2b::Flpo*(*32*); *Phox2b::Cre*(*58*), *vGlut2::Cre*(*59*), *Atoh1::Cre* (*13*), *Atoh1::FRTCre* (*13*), *Olig3::Cre^ERT2^* (*31*), *Foxg1^iresCre^*, *RC::FELA*(*60*) *Tau::Syp-GFP* (*61*), *Rosa::nlsLacZ* (also known as *Tau^mGFP^*)(*62*) and *Ai9* (*63*). For behavioral experiments, all mice were produced in a B6D2 background.

The *Rosa^FTLG^* mutant mouse line was established at the Institut Clinique de la Souris (Phenomin-ICS), Illkirch, France). The targeting vector was constructed as follows. A PCR fragment containing the rat synaptophysin cDNA fused to GFP was cloned by SLIC cloning with a 346 bps double stranded synthetic HSV TK pA followed by an 29 bps homology for the 5’ extremity of the 3’ Rosa homology arm plus an NsiI site, in an ICS proprietary vector containing a floxed NeoR-STOP cassette. In the second cloning step, the NeoR cassette was removed by BamHI and SpeI restriction digests and replaced by SLIC cloning with the cDNA of tdTomato. The third cloning step introduced, 5’ of the floxed tdTomato-STOP cassette, a DNA fragment containing an *NsiI* site followed by a 29 bps homology for the 3’ of the pCAG, followed by a MCS. The fourth step was the cloning of a FRT-surrounded NeoR-STOP cassette previously excised from an ICS proprietary vector in the *SmaI* site of the restriction site introduced in the MCS cassette. Finally, a fifth cloning step comprised the excision of a 7.8 kb fragment containing the whole FRT-NeoR-STOP-FRT LoxP-TdTomato-STOP-LoxP Syn-YFP cassette by a *NsiI* digest and its subcloning via SLIC cloning in an ICS proprietary vector containing a pCAG (Chicken b-actin promoter preceded by a CMV enhancer) and both 5’ and 3’ Rosa homology arms. The linearized construct was electroporated in C57BL/6N mouse embryonic stem (ES) cells (ICS proprietary line). After G418 selection, targeted clones were identified by long-range PCR and further confirmed by Southern blot with an internal (Neo) probe and a 5’ external probe. One positive ES cell clone was validated by karyotype spreading and microinjected into BALB/c blastocysts. Resulting male chimeras were bred with wild type C57BL/6N females. Germline transmission was achieved in the first litter.

#### Housing

Animals were group-housed with free access to food and water in controlled temperature conditions and exposed to a conventional 12-h light/dark cycle. Experiments were performed on embryos at embryonic (E) days E11.5-17.5, neonate pups at postnatal day 2-8 (P2-8) and adult (P30-56) animals of either sex. All procedures were approved by the French Ethical Committee (authorization 26763-2020022718161012) and conducted in accordance with EU Directive 2010/63/EU. All efforts were made to reduce animal suffering and minimize the number of animals.

### Viral vectors for tracing, optogenetic and photometry experiments

For anterograde tracing from Peri5^*Atoh1*^ and IRT^*Phox2b*^ we injected unilaterally 250 nl of a Cre-dependent AAV2/8-hSyn-FLEX-mGFP-2A-Synaptophysin-mRuby (Titer: 1.3×10^12^ vg/ml, Viral Core Facility Charité).

For retrograde trans-synaptic tracing from muscles we injected unilaterally 50 to 100 nl of a 1:1 viral cocktail comprised of RV-B19-ΔG-mCherry or RV-B19-ΔG-GFP (titer: 1.3×10^9^ and 5.8×10^8^ TU/ml respectively, Viral Vector Core - Salk Institute for Biological studies) and a HSV-hCMV-YFP-TVA-B19G (titer: 3×10^8^ TU/ml, Viral Core MIT McGovern Institute).

For retrograde tracing from IRT^*Phox2b*^ we injected unilaterally 250 nl of a Cre-dependent AAV1/2-Syn-flex-nGToG-WPRE3 (Titer: 8.1×10^11^ viral genomes (vg)/ml, Viral Core Facility Charité). Two weeks later we injected EnvA-RV-B19-ΔG-mCherry (Titer: 3.1×10^8^ vg/ml, Viral Vector Core, Salk Institute for Biological studies).

For optogenetic and photometry experiments we respectively injected 250 nl of AAV1/2-Ef1a-DIO-stCoChR-P2A-mScarlet (titer: 3×10^13^ vg/ml, kind gift from O. Yzhar) or 250 nl of AAV1-syn-FLEX-jGCaMP7s-WPRE (titer: 1×10^12^ vg/ml Addgene #104487-AAV1).

### Surgical procedures

#### Stereotaxic injections and implants

All surgeries were conducted under aseptic conditions using a small animal digital stereotaxic instrument (David Kopf Instruments). Mice were anesthetized with isoflurane (3.5% at 1 l/min for induction and 2-3% at 0.3 l/min for maintenance). Buprenorphine (0.025 mg/kg) was administered subcutaneously for analgesia before surgery. A feed-back-controlled heating pad was used to maintain the animal temperature at 36°C. Anesthetized animals were placed in a stereotaxic frame (Kopf), a 100 μl injection of lidocaine (2%) was made under the skin covering the skull, after which a small incision was made in the scalp and burr-free holes were drilled in the skull to expose the brain surface at the appropriate stereotaxic coordinates [anterior-posterior (AP) and medial-lateral (ML) relative to bregma; dorsal-ventral (DV) relative to brain surface at coordinate (in mm)]: −4.9 AP, 1. 2 ML, 4.0 DV to target the Peri5^*Atoh1*^ neurons; −6.7 AP, 0.5 ML, 4.2 DV to target the IRT^*Phox2b*^ neurons. A 0.5 ML coordinate was selected for virus deliveries to the IRt^*Phox2b*^ to circumvent the potential infection of nTS neurons along the injecting pipette track, a 4.0 DV coordinate was selected for virus deliveries to the Peri5^*Atoh1*^ to target the center of Mo5. Viral vectors were delivered using glass micropipettes (tip diameter ca. 100mm) backfilled with mineral oil connected to a pump (Legato 130, KD Scientific, Phymep, France) via a custom-made plunger (Phymep, France). The injector tip was lowered an additional 0.1 mm below the target site and then raised back to the target coordinate before infusion started (flow of 25 nl/min) to restrict virus diffusion to the site of injection and prevent leakages along the needle track. After infusion, the injection pipette was maintained in position for 10 minutes, then raised by 100 μm increments to retract the pipet from the brain. For optogenetic and photometry experiments, 200 μm core optic fibers (0.39 NA and 0.57 NA respectively) (Smart Laser Co., Ltd) were implanted following vector injections, ~500μm above the sites of interest (−4.9 AP, 1.2 ML, 3.0 DV for Peri5^*Atoh1*^; −6.7 AP, 0.9 ML, 3.6 DV for IRT^*Phox2b*^. The optic fibers were secured via a ceramic ferrule to the skull by light-cured dental adhesive cement (Tetric Evoflow, Ivoclar Vivadent). Mice recovered from anesthesia on a heating pad before being placed, and monitored daily, in individual cages.

#### Intramuscular injections

All surgeries were conducted under aseptic conditions on P2 neonates anesthetized by deep hypothermia. For induction, pups were placed in latex sleeves gently buried in crushed ice for 3-5 minutes and maintenance (up to 15 min) was achieved by placing anesthetized pups on a cold pack (3-4°C). Following small incisions of the skin to expose the targeted muscles, 0.5 μl of the viral cocktail (or 0.5% Cholera toxin subunit B (CTB) (List Labs) for labeling of the infrahyoids) were injected via a pneumatic dispense system (Picospritzer) connected to a glass pipette (tip diameter ca. 0.1 mm) mounted on a 3D micromanipulator to guide insertion in the desired muscle. Typically, 5-10 pressure pulses (100 ms, 3-5 bars) were delivered while muscular filling was checked visually by the spreading of Fast-Green (0.025%) added to the viral solution. The pipette was withdrawn and the incision irrigated with physiological saline and closed using 10-0 gage suture (Ethilon). The mouse was placed on a heat pad for recovery and returned to the mother. Six days post-injection (4 days for CTB), pups were deeply anesthetized, transcardially perfused with 4% paraformaldehyde (PFA) in phosphate-buffered saline (PBS), and the brains was dissected out and post-fixed overnight in 4% PFA, and cryoprotected in 15% sucrose in PBS and were stored at −80°C.

### Histology

#### Immunofluorescence

Depending on the stage, the brain was analyzed in whole embryos dissected out of the uterine horns up to E16.5, dissected out from decapitated embryos from E17.5 to P0 or, after P0, dissected in cold PBS from euthanized animals perfused with cold PBS followed by 4% paraformaldehyde. Brains or embryos were post-fixed in 4% paraformaldehyde overnight at 4 °C, rinsed in PBS and cryoprotected in 15% sucrose overnight at 4°C. Tissues were then frozen in Tissue-Tek® O.C.T. compound for cryo-sectioning (14-30 μm) on a CM3050s cryostat (Leica). Sections were washed for 1 hour in PBS and incubated in blocking solution (5% calf serum in 0.5% Triton-X100 PBS) containing the primary antibody, applied to the surface of each slide (300 μl per slide) placed in a humidified chamber on a rotating platform. Incubation was for 4-8 hours at room temperature followed by 4°C overnight. Sections were washed in PBS (3 × 10 minutes), then incubated with the secondary antibody in blocking solution for 2 hours at room temperature, then washed in PBS (3 × 10 minutes), air-dried, and mounted under a cover slip with fluorescence-mounting medium (Dako). Primary antibodies used were: goat anti-Phox2b (RD system AF4940, diluted 1:100), rabbit anti-peripherin (Abcam ab4666, 1:1000), guinea-pig anti-Lmx1b (Müller et al.,2002, 1:1000), goat anti-ChAT (Millipore AB144p), 1:100), chicken anti-βGal (Abcam, ab9361, 1:1000), chicken anti-GFP (Aves labs, GFP-1020,1:1000), goat anti-ChAT (Millipore, AB144p, 1:100), rabbit anti-GFP (Invitrogen, A11122, 1:1000), rabbit anti-Phox2b (Pattyn et al., 1997,1:500), rat anti-RFP (Chromotek, 5F8, 1:1000), goat anti-CTB (List Labs, #703, 1:500). All secondary antibodies were used at 1:500 dilution: donkey anti-chicken 488 (Jackson laboratories, 703-545-155), donkey anti-chicken Cy5 (Jackson laboratories, 703-176-155), donkey anti-goat Cy5 (Jackson laboratories, 705-606-147), donkey anti-rabbit 488 (Jackson laboratories, 711-545-152) donkey anti-rabbit Cy5 (Jackson laboratories, 712-165-153), donkey anti-rat Cy3 (Jackson laboratories, 711-495-152), donkey anti-Guinea pig Cy3(706-165-148). Epifluorescence images were acquired with a NanoZoomer S210 digital slide scanner (Hamamatsu Photonics) and confocal images with a Leica SP5 confocal microscope (Leica). Pseudocoloring, image brightness and contrast were adjusted using Adobe Photoshop and ImageJ.

#### In Situ Hybridization and immunohistochemistry

For the *Atoh1* probe, primers containing SP6 and T7 overhangs were used to amplify a 607 bp region from a plasmid containing the full length *Atoh1* CDS. The purified amplicon was then used as the template for antisense probe synthesis with T7 RNA polymerase using the following primers: Forward Primer: 5’-CGATTTTAGGTGACACTATAGAAATCAA-CGCTCTGTCGGAGTT-3’; Reverse Primer: 5’-CTAATACGACTCACTATAGGGACAGAGGAAGGGGAT-TGGAAGAG −3’. To generate the *Cited1* probe, a 687 bp fragment of the murine *Cited1* gene was amplified from E13.5 mouse brain cDNA (superscript III kit, Invitrogen) and cloned into pGEM-T vector (Promega), using the following primers: Forward Primer: 5’-TGGGGGGCTTAAGAGCCCGG-3’; Reverse Primer: 5’-AGGTGAGGGGTAGGATGCAG-3. pGEM clones were linearized with NotI and transcribed with SP6 or T7 RNA polymerase using the DIG RNA labelling Kit (Roche 1277073) to generate antisense or sense probes. In situ hybridization was performed on 14 μm thick cryo-sections. Sections were washed for 10 min in PBS prepared in DEPC-treated water, then washed in RIPA buffer (150 mm NaCl, 1% NP-40, 0.5% Na-deoxycholate, 0.1% SDS, 1 mM EDTA, 50 mM Tris, pH 8.0) for 20 min, post fixed in 4% paraformaldehyde for 15 min followed by rinses in PBS (3 × 10 min). Whenever ISH was to be followed by an immunohistochemical reaction, slides were incubated for 30 minutes in a mixture of 100% ethanol and 0.5% H_2_0_2_, washed in PBS (3 × 10 mins), then incubated in Triethanolamine containing 0.25% acetic acid for 15 minutes and washed again in PBS (3 × 10 mins). Antisense RNA probes were diluted in 200μl hybridization buffer (5 × SSC, 10% dextran sulfate, 500μg/mL Herring sperm DNA, 250μg/mL Yeast-RNA, 50% formamide) and denatured at 95°C for 5 minutes, cooled briefly on ice, then diluted at 100-200ng/ml in 17ml hybridization buffer for incubation in slide mailers, at 70°C overnight. The next day, slides were washed for 1 hour at 70°C in 2 × SSC, 50% formamide and 0.1% Tween 20 and for 1 hour in 0.2 × SSC at 70°C. Slides were washed in B1 buffer (0.1M Maleic acid; pH 7.5, 0.15M NaCl, 0.1% Tween 20), 3 X10 min. The sections were then blocked for 1 h at room temperature by incubation in blocking buffer (B1 buffer supplemented with 10% heat-inactivated fetal calf serum). The blocking solution was replaced by alkaline phosphatase-conjugated anti-DIG antibody (Roche diagnostics, 11093274910) diluted 1:200 in the blocking buffer and sections were incubated overnight at 4°C under cover slip. The following day the slides were rinsed in B1 buffer (3 × 10 min), equilibrated with B3 buffer (0.1 M Tris pH 9.5, 0.1 M NaCl, 50 mM MgCl_2_, 0.1% Tween 20) for 30 min and colorimetric detection of the digoxigenin-labeled probe was performed with NBT-BCIP substrate for alkaline phosphatase (Thermo Scientific). The reaction was stopped by washing the slides in PBS-0.1% Tween 20 (2 × 5 min) and fixing in 4% paraformaldehyde for 15 min. Sections were then washed in PBS-0.1% Tween 20 for 5 min each. Sections were incubated in blocking buffer (10% fetal calf serum diluted in 0.1% Tween 20 in PBS) for 1 hour at room temperature, then in blocking buffer containing the primary antibody at 4°C overnight. The next day, slides were washed for 10 min and biotinylated secondary antibody (diluted at 1:200 in blocking buffer) was applied for 2h at room temperature and peroxidase enzyme detection of biotinylated antibody was carried out as per manufacturer’s guidelines with the Vectastain Elite ABC kits (PK-6101 and PK-4005; Vector Laboratories), followed by color development using 3, 3’-Diaminobenzidine (SIGMA FAST D4293-50SET). The reaction was stopped by washing the slides for 2 × 5 min in Milli-Q water, then sections were allowed to air-dry completely before mounting with Aquatex (Sigma Aldrich) for microscopy. Hybridized sections were imaged with a Leica DFC420C camera mounted on a Leica DM5500B microscope.

### Data Analysis of histology

#### Counts of premotor neurons and Lmx1b neurons

Cells expressing *mCherry* and/or *nlsLacZ*, were counted in a spheroid of fixed dimension and position delimitating the ipsilateral dorsal IRt, drawn on the approximately 7 sections that were in register with the compact formation of MoA; n=4 animals, 87±20 SEM premotor neurons per animal.

Cells expressing *Phox2b* and/or *Lmx1b* were counted as above from one side; n=3 animals, 1321±46 SEM neurons per animal.

#### Inputs to IRt^Phox2b^

Images of sections were aligned to the Allen Brain Atlas using QuickNII (*64*) (https://www.nitrc.org/projects/quicknii). Labelled neurons were manually annotated as IRt seed neurons (GFP^+^ mCherry^+^) or monosynaptic input neurons (mCherry^+^) in ImageJ. The pixel coordinates of identified input neurons were transformed into Allen Brain Atlas coordinates as previously described (*65*)(*66*) and corresponding Allen Brain Atlas brain structures identified using CellfHelp (https://github.com/PolarBean/CellfHelp). Data from individual replicates were tabulated, normalized and pooled to generate a list of brain regions that provide monosynaptic input to IRt^*Phox2b*^. The display bar graph excludes any input below 0.3%.

### Behavioral experiments

#### Timing and training

All behavioral experiments started four weeks after the viral injection. Two weeks after surgery animals were habituated to head-fixation through sessions of increasing duration (2 minutes) every other day, starting at 2 minutes on day 0 and a final duration of 10 minutes on day 5 which corresponded to the duration of recording sessions. Animals were given condensed milk as a reward after each session. Animals used for photometry experiments were introduced to a lick port during habituation. During acquisition or manipulation animals were head-fixed within a 5cm tube, illuminated from below and above by an LED light. Animals were water deprived for 12 hours prior to photometry experiments.

#### Optogenetics

For optogenetic photostimulation of st-CoChR expressing neurons, fiber optic canulae were connected to a 473-nm DPSS laser (CNI, Changchun, China) through a patch cable (200 μm, 0.37 NA) and a zirconia mating sleeve (Thorlabs). Laser output was controlled using a pulse generator (accupulser, WPI), which delivered single continuous light pulses of 50-1000 ms. Light output through the optical fibers was adjusted to ~5 mW at the fiber tip using a digital power meter (PM100USB, Thorlabs), to prevent heat. All light stimuli were separated by minimal periods of 10s. Laser output was digitized at 1kHz by a NI USB-6008 card (National Instruments) and acquired using a custom-written software package (Elphy by G Saddoc, https://www.unic.cnrs.fr/software.html).

#### Photometry

For photometry experiments, a single site fiber photometry system (Doric Lenses Inc, Canada) was used to measure the excited isobestic (405nm) and calcium-dependent fluorescence of jGCaMP7s (465nm). Doric neuroscience studio software system (Doric Lenses Inc, Canada) was used to operate the photometry hardware and acquire the photometry signal. Briefly, using the “lock in mode” function, 465 nm and 405nm LEDs were sinusoidally modulated at 208.616 Hz and 572.205 Hz respectfully (to avoid any electrical system harmonics at 50/60 Hz, 100/120 Hz, 200/240 Hz) at an intensity of 30 μW and coupled to a patch cable (diam. 200μm, 0.57nA) after passing through an optical assembly (iLFMC4, Doric Lenses Inc, Canada). The modulated excitation signal was then directed through an implanted fiber optic cannula (diam. 200μm, 0.57NA) onto the IRt via the mated patch cable and the emitted signal was then returned via the same patch cable to a fluorescence detector head, mounted on the optical assembly and amplified. The raw detected signal was acquired at 12 kHz and then demodulated in real time to reconstitute the excited isobestic (405nm) and calcium dependent GCaMP (465nm) signals. Contact between the tongue and the lick port during spontaneous licking bouts were registered via a SEN-1204 capacitance sensor (Sparkfun) connected to Arduino Uno R3 microcontroller board (Arduino) and acquired at 12 kHz via the Doric fibre photometry console.

#### Automated markless pose estimation

Spontaneous and light-evoked licking sequences were filmed at portrait (Fig. 4A) and profile angles (Fig. 4D) with a CMOS camera (Jai GO-2400-C-USB) synchronized by a 5V TTL pulse. The acquired frames (800 × 800 pixels, 120 fps,) were streamed to a hard disk using 2ndlook software (IO Industries) and compressed using an MPEG-4 codec. Portrait views were used for video tracking of optogenetically-evoked oromotor movements, while profile views were preferably used for photometry experiments, to optimize detection of the tongue, which was partially obscured by the nearby lick port when filmed from the portrait angle.

Using DeepLabCut (version 2.0.7,(*67*)), we trained 2 ResNet-50 based neural networks to identify the tip of the tongue and lower jaw from portrait and landscape views (**Fig. 4 A,B,D**). The “portrait” network was trained on a set of 264 frames (800 × 800 pixels) derived from 11 videos of 6 different mice for >400,000 iterations, reporting a train error of 1.85 pixels and test error of 6.79 pixels upon evaluation. The “profile” network was trained on a set of 90 frames (800 × 800 pixels) from 4 videos of 4 different mice for >800,000 iterations reporting a train error of 1.66 pixels and test error of 4.57 pixels upon evaluation. These networks were then used to generate Cartesian estimates for the Y-axis position of the jaw and tongue for experimental videos.

### Data analysis

We analyzed behavioral and fiber photometry data using custom written Python scripts (Python version 3.7, Python Software Foundation). In most instances, mice underwent multiple sessions of the same experiment. These sessions were then averaged and treated as a single replicate for that animal. Fiber photometry and photostimulation data was resampled to 120 Hz to match the acquisition rate of video recordings.

#### Fiber photometry

Photometry data was analyzed as previously described (*44*). Both 465 nm and 405 nm signals were first low-pass filtered. The 465 nm signals was then normalized using the function ΔF/F = (F − F0) / F0, in which F is the 465 nm signal, and F0 is the least-squared mean fit of the 405 nm signal. Responses (photometry, jaw and tongue) were then aligned to the peak of the first derivative of jaw opening events that were active during a lick bout.

#### Statistical analysis of fiber photometry data

For each recording session in one animal, correlations between lick port contact and calcium signals were computed for all possible shifts at 120Hz spanning from −10s to +10s, producing one curve per session (Fig 3G). A null correlation curve per recording session was constructed by performing the same computation after shuffling the lick port contact (Fig 3G). All recording sessions and all null correlation curves were averaged for each animal, to produce a single mean shifted correlation curve and a null mean correlation curve per animal (Fig. S4). The maxima values of both shifted and null mean curves were retrieved for each animal (n=4). A paired t-test between these values indicated a shifted correlation between both signals.

#### Normalisation of jaw and tongue pose estimation

Cartesian pixel estimates of the jaw and tongue were corrected to a 5 mm scale bar within the video frame, and smoothed using the Savitzky-Golay filter. For optogenetic experiments, the jaw position was normalized to its averaged location 50-100 ms prior to stimulation. For fiber photometry experiments, the jaw position was normalized to its average location during quiescent periods between 1-3 s long. As the tongue was only present during stimulation of IRt^*Phox2b*^ or spontaneous lapping, we normalized the tongue distance empirically by observing the first detected instance of tongue protrusion that succeeded jaw-opening events. All positional estimates of the tongue that had a probability < 5 % (*67*) were then set to the empirically determined baseline to filter out aberrant estimates of the tongue position during periods of the recording where it was not visible. Fiber photometry

#### Licking frequency

For data collected during optogenetic experiments we first obtained the onsets of each lick during a 1000 ms stimulation window. These onsets were identified as the peaks of the first derivate of each lick within a lick bout. Lick frequency was then calculated as the number of lick events divided by length of time from the last lick to the first lick within a lick bout. For data collected during fiber photometry, lick frequency was determined by the number of contact events of the capacitance sensor divided by the length of time from the last to the first lick within a lick bout.

#### Statistical Analysis

All data are reported as mean ± s.e.m (shaded area). P values for independent samples comparison were performed using a two-tailed Student’s t-test.

## Acknowledgments

We thank the animal facility of IBENS, the imaging facility of IBENS (supported by grants from Fédération pour la Recherche sur le Cerveau, Région Ile-de-France DIM NeRF (2009 and 2011) and France-BioImaging), Ofer Yitzar for the *AAV1-EF1a-DIO-stCoChR-P2A-mScarlet* vector. The mouse *Rosa^FTLG^* mutant line was established at the Institut Clinique de la Souris (Phenomin-ICS) in the Genetic Engineering and Model Validation Department. Funding is from CNRS, École Normale Supérieure, INSERM, Association Nationale pour la Recherche ANR - 15-CE16-0013 (to JFB), Association Nationale pour la Recherche ANR-17-CE16-0006 (to JFB) and ANR-19-CE16-0029 (to GF), Fondation pour la Recherche Médicale DEQ2000326472 (to JFB), ‘Investissements d’Avenir’ program ANR-10-LABX-54 MEMO LIFE and ANR-11-IDEX-0001–02 PSL Research University), and Région Ile-de-France (to SS).

## Author contributions

Conceptualization: BD, CG, GF, JFB

Investigation: BD, SS, PB, ERH, ZC, SD, SA

Formal Analysis: SM, HC, AG

Supervision: CG, GF, JFB, JFAP

Resources CB

Writing BD, GF, JFB

## Data and materials availability

All data needed to evaluate the conclusions of the paper are available in the main text or the supplementary materials, or from the corresponding authors upon reasonable request.

## Supplementary figures

**Fig. S1.**
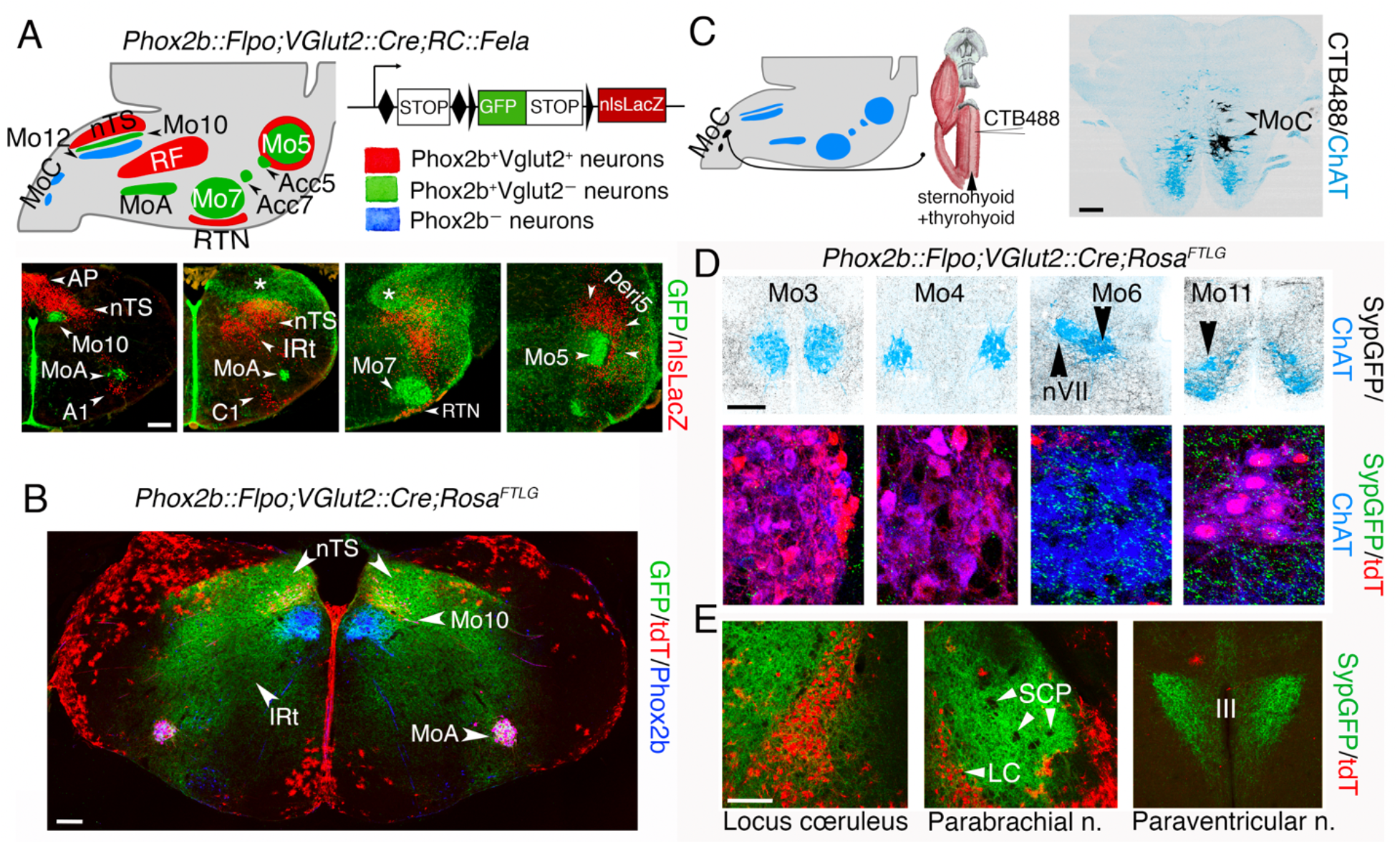
**(A)** (Upper) Strategy for transgenic labeling of glutamatergic Phox2b^+^ interneurons using the *FELA* transgene, and summary of the results. (Lower) Coronal sections through the pons and medulla of a *vGlut2::Cre;Phox2b::Flpo;RC::Fela* embryo at E17.5. Branchiomotor and visceromotor neurons (cholinergic and Phox2b^+^) express GFP (green), while interneurons that are glutamatergic and Phox2b^+^ express *nlsLacZ* (red). The dorsal aspect of the hindbrain harbors numerous GFP+ astrocytes (asterisks), likely born from Phox2b^+^ progenitors, in p3/pMNv (*12*)(*13*) or possibly dB2. Coronal section through the medulla of a *vGlut2::Cre;Phox2b::Flpo;Rosa^FTLG^* embryo at E17.5, showing that branchial and visceral neurons have retained tdT expression (since they undergo recombination by *Flpo* alone), while glutamatergic *Phox2b^+^* neurons in the nTS or IRt have lost tdT, to gain *SypGFP* expression after dual *Cre* and *Flpo* recombination. *Syp-GFP* marks fields of synaptic boutons in discrete areas of the medulla. (**C**) (left) Strategy for labeling the motor nucleus for the infrahyoid muscles (sternohyoid and thyrohyoid) (MoC), by retrograde transport of CTB. (right) transverse section at the spinal-medulla junction showing two islands of labelled motoneurons (black). **(D)** Motor nuclei for extraocular muscles (Mo3, Mo4 and Mo6) and for the trapezius and sternocleidomastoid (Mo11), do not receive labelled boutons in a *vGglut2::Cre;Phox2b::Flpo;Rosa^FTLG^* background at P8. (**E**) Known sites of projection from Phox2b^+^ glutamatergic interneurons (including C1 neurons and the nTS) are covered by GFP+ boutons: the parabrachial nucleus (*14*), the locus coeruleus and the paraventricular nucleus of the hypothalamus (*15*). LC, locus coeruleus; SCP, superior cerebellar peduncle; III, third ventricle. Scale bars, 200 μm.

**Fig. S2.**
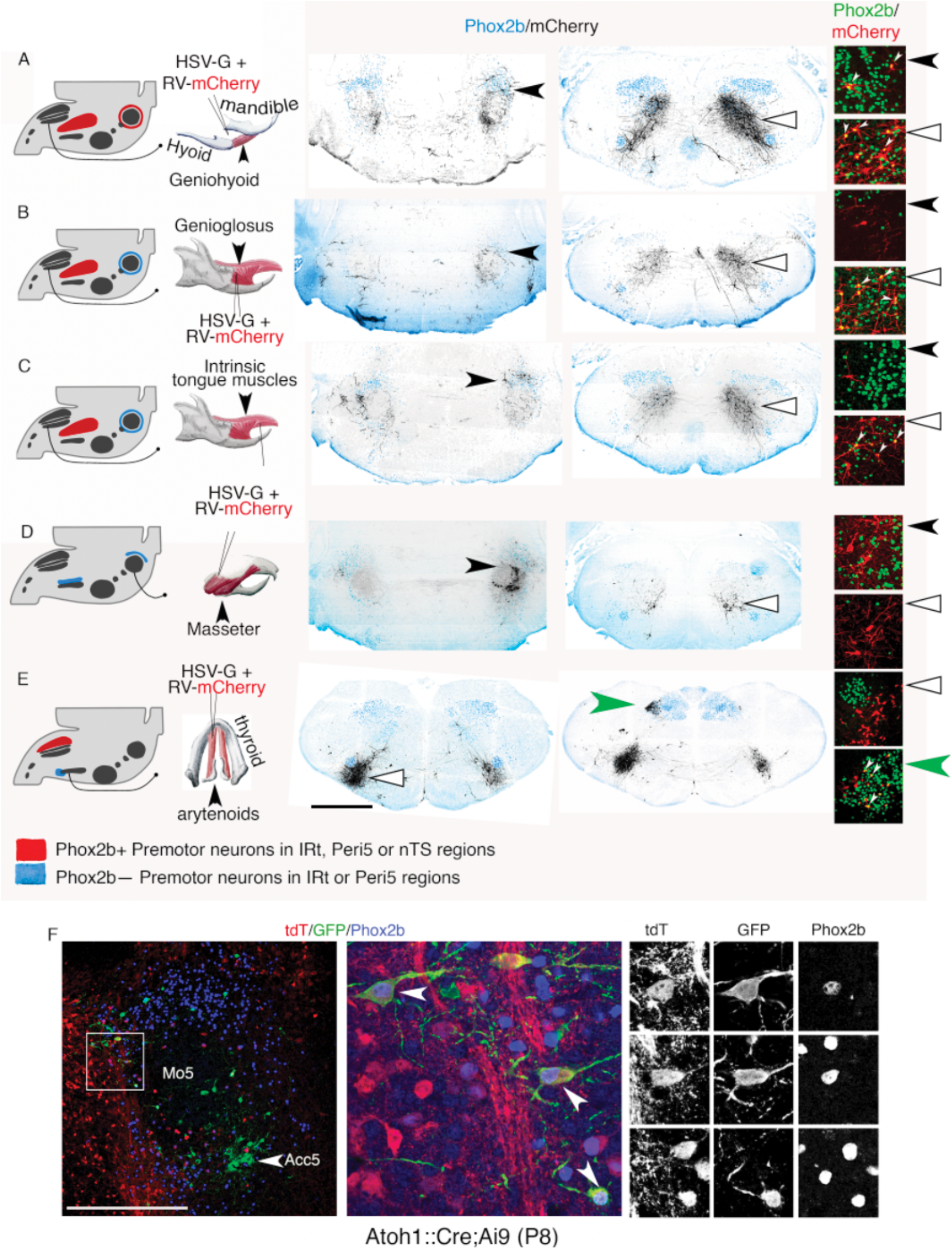
Retrograde monosynaptic labeling of the premotor neurons for the geniohyoid. (**A**), genioglossus (**B**), intrinsic tongue muscles (**C**), masseter (**D**) and thyro-arytenoid (**E**). (Left column) Labeling strategy and summary of the results. (Middle column) Sections through the pons at the level of Mo5 (**A**-**D**, left), or at the level of the IRt (**A**-**D**, right, and **E**, left and right) showing the filled premotor neurons (black) on the landscape of Phox2b+ neurons (blue). (Right column) Close ups of filled premotor neurons (red) either expressing *Phox2b* (green nuclei), or not. The thyro-arytenoid and masseter have no Phox2b^+^ premotor neurons in the IRt or Peri5. (**F**) Coronal section through the peri5 region of a Cre-reporter *Ai9* mouse crossed with *Atoh1::Cre*, whose posterior digastric muscle was co-injected with a DG-rabies virus encoding GFP and a helper HSV-G, counter-stained for *Phox2b*, at 3 magnifications from left to right. Three triple-labeled neurons are highlighted, which thus have a history of both *Atoh1* and *Phox2b* expression, and are premotor to the posterior digastric. Scale bar, (**A**-**E**), 1 mm; (**F**), 500μm.

**Fig. S3.**
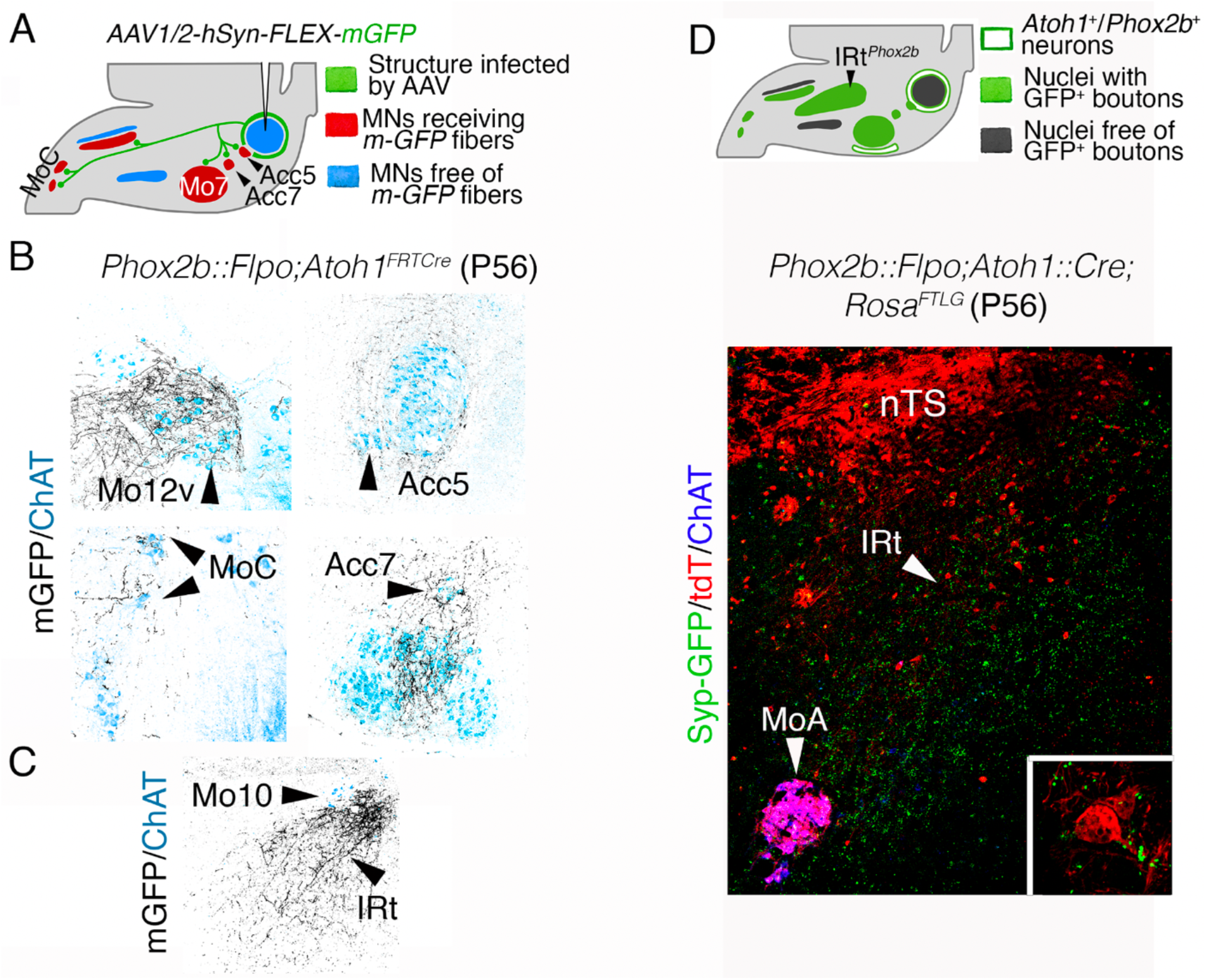
(**A**): Strategy for viral tracing the projections of Peri5^*Atoh1*^ and summary of the results. (**B)**: Coronal sections through the motor nuclei (ChAT^+^, blue) that receive projections from Peri5^*Atoh1*^, labelled with *mGFP* encoded by the AAV anterograde virus (black). (**C**): Coronal sections through the medulla showing *mGFP-*labeled fibers in the IRt. (**D**): Strategy for transgenic labeling projections from *Atoh1^+^/Phox2b^+^* cells (Peri5^*Atoh1*^ and RTN) using the *Rosa^FTLG^* transgene and summary of the results already visible in Fig 3A. (**D**) Projections from Peri5^*Atoh1*^ +RTN on IRt^*Phox2b*^ at intermediate and high (inset) magnifications.

**Fig. S4.**
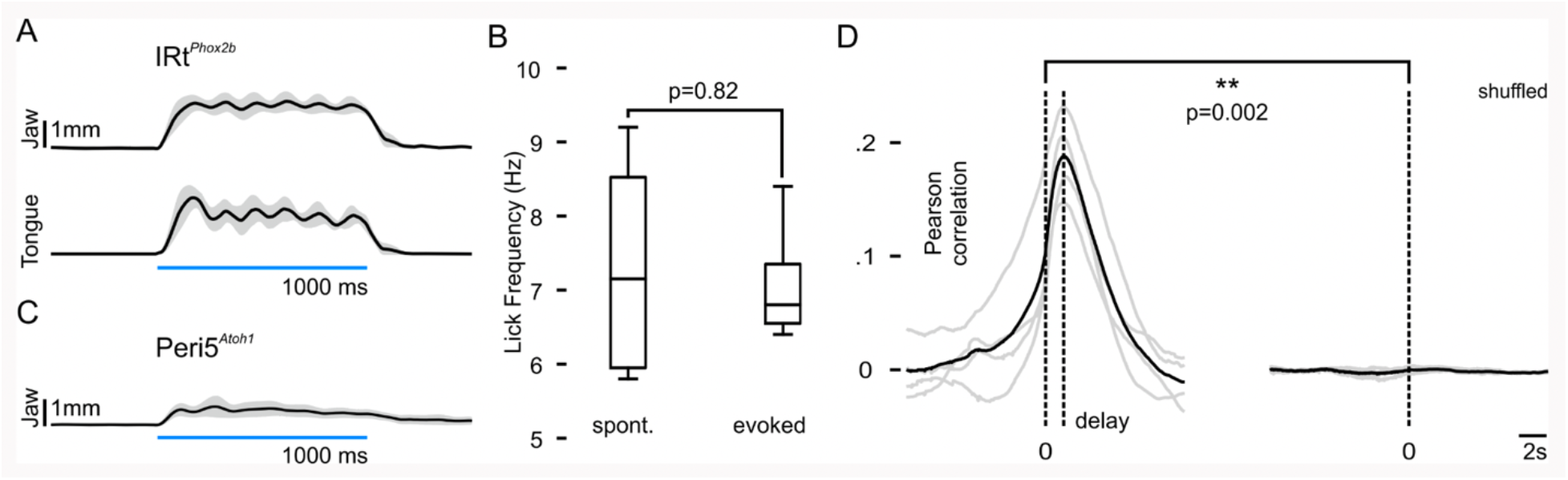
(**A**) Grand average of tracked position of jaw and tongue on the Y axis upon 1000 ms stimulation of IRt^*Phox2b*^ (n=4 mice, 34 trials). (**B**) Box plot of spontaneous (7.3 Hz ±0.8 SEM) (n=4) and evoked licking frequency (7.1 Hz ±0.4 SEM). (**C**) Grand average (n=4 mice, 21 trials) of tracked position of the tongue on the Y axis upon 1000 ms stimulation of Peri5^*Atoh1*^. (**D**) (left) Mean shifted correlation curves displayed for each animal (n=4, gray) and the overall mean (black), displaying a 1.2s delay between lick port contact and maximum correlation; (right) same computation on the same data but with the lick signal shuffled.

**Movie S1.**

Stimulation (1000 ms) of IRt^*Phox2b*^. Displayed at ¼ speed.

**Movie S2.**

Stimulation (1000 ms) of Peri5^*Atoh1*^. Displayed at ¼ speed

**Movie S3.**

Activity of IRt^*Phox2b*^ during a spontaneous licking bout. Displayed at ¼ speed.

